# Molecular basis of RanGTP-activated nucleosome assembly with Histones H2A-H2B bound to Importin-9

**DOI:** 10.1101/2023.01.27.525896

**Authors:** Joy M. Shaffer, Jenny Jiou, Kiran Tripathi, Oladimeji S. Olaluwoye, Ho Yee Joyce Fung, Yuh Min Chook, Sheena D’Arcy

## Abstract

Padavannil et al. 2019 show that Importin-9 (Imp9) transports Histones H2A-H2B from the cytoplasm to the nucleus using a non-canonical mechanism whereby binding of a GTP-bound Ran GTPase (RanGTP) fails to evict the H2A-H2B cargo. Instead, a stable complex forms, comprised of equimolar RanGTP, Imp9, and H2A-H2B. Unlike the binary Imp9•H2A-H2B complex, this RanGTP•Imp9•H2A-H2B ternary complex can release H2A-H2B to an assembling nucleosome. Here, we define the molecular basis for this RanGTP-activated nucleosome assembly by Imp9. We use hydrogen-deuterium exchange coupled with mass spectrometry and compare the dynamics and interfaces of the RanGTP•Imp9•H2A-H2B ternary complex to those in the Imp9•H2A-H2B or Imp9•RanGTP binary complexes. Our data are consistent with the Imp9•H2A-H2B structure by Padavannil et al. 2019 showing that Imp9 HEAT repeats 4-5 and 18-19 contact H2A-H2B, as well as many homologous importin•RanGTP structures showing that importin HEAT repeats 1 and 3, and the h8 loop, contact RanGTP. We show that Imp9 stabilizes H2A-H2B beyond the direct binding site, similar to other histone chaperones. Importantly, we reveal that binding of RanGTP releases H2A-H2B interaction at Imp9 HEAT repeats 4-5, but not 18-19. This exposes DNA- and histone-binding surfaces of H2A-H2B, thereby facilitating nucleosome assembly. We also reveal that RanGTP has a weaker affinity for Imp9 when H2A-H2B is bound. This may ensure that H2A-H2B is only released in high RanGTP concentrations near chromatin. We delineate the molecular link between the nuclear import of H2A-H2B and its deposition into chromatin by Imp9.

**Significance:** Imp9 is the primary importin for shuttling H2A-H2B from the cytoplasm to the nucleus. It employs an unusual mechanism where the binding of RanGTP alone is insufficient to release H2A-H2B. The resulting stable RanGTP•Imp9•H2A-H2B complex gains nucleosome assembly activity as H2A-H2B can be deposited onto an assembling nucleosome. We show that H2A-H2B is allosterically stabilized via interactions with both N- and C-terminal portions of Imp9, reinforcing its chaperone-like behavior. RanGTP binding causes H2A-H2B release from the N-terminal portion of Imp9 only. The newly-exposed H2A-H2B surfaces can interact with DNA or H3-H4 in nucleosome assembly. Imp9 thus plays a multi-faceted role in histone import, storage, and deposition regulated by RanGTP, controlling histone supply in the nucleus and to chromatin.

## Introduction

In the nucleus, 147-bp segments of DNA wrap around an octamer of core histones to form nucleosomes (1). The histone octamer comprises two copies of histones H2A-H2B and H3-H4 that are synthesized and folded in the cytoplasm, and then actively transported to the nucleus (2, 3). Nuclear import receptors (importins) of the Karyopherin-β family play a central role in chromatin regulation through this histone transport process. Cellular and proteomic studies show that H2A-H2B is primarily imported by yeast Kap114 and its human paralog Importin-9 (Imp9, also known as IPO9) (3–5). Kap114 and Imp9 have 21.2% sequence identity and share a super-helical HEAT-repeat fold containing 20 repeats (h1-h20), each with antiparallel helices a and b (6, 7). Imp9 and Kap114 also contain three loop insertions; the h8 loop, the h18-h19 loop, and an acidic h19 loop (6, 7). Previous studies have shown that Imp9 has histone chaperone activity and prevents H2A-H2B from spuriously interacting with other macromolecules, especially DNA (8). The crystal structure of Imp9 with H2A-H2B shows contacts between the N-terminal (h2-h5) and C-terminal (h17-h19) repeats of Imp9 with opposite ends of H2A-H2B (6). These extensive interactions block H2A-H2B surfaces that bind histones and DNA in the nucleosome.

The binding between importins and their cargo is regulated by Ran GTPases (9–17). Most Ran in the cytoplasm is loaded with GDP, while nuclear Ran is loaded with GTP (RanGTP), with the concentration gradient between these two forms driving nucleo-cytoplasmic transport (18, 19). This gradient is established by chromatin-binding protein, Ran guanine nucleotide exchange factor (RCC1), which catalyzes the exchange of GDP to GTP in nuclear Ran; and GTPase Activating Protein (RanGAP) that catalyzes the exchange of GTP to GDP in cytoplasmic Ran (17, 20, 21). The binding between an importin and RanGTP is highly conserved in paralogs and orthologs with the N-terminal HEAT repeats (h1-h4) of the importin contacting Switch 2 of the RanGTP. The importin repeats that contact Switch 1 of RanGTP are Karyopherin specific (10, 17, 22–26). In most nuclear import systems, the binding of RanGTP to the importin causes cargo dissociation such that the cargo-free importin then travels back to the cytoplasm (9–17). The complex of Imp9 and H2A-H2B is atypical as the binding of RanGTP is insufficient to dissociate the H2A-H2B cargo, and a stable RanGTP•Imp9•H2A-H2B complex forms (6). When RanGTP binds, however, it does facilitate the transfer of H2A-H2B to a nucleosome assembly intermediate (6). Before this work, the molecular details of the ternary RanGTP•Imp9•H2A-H2B complex and the mechanism of RanGTP-induced H2A-H2B nucleosome deposition were unknown.

Here, we uncover the molecular basis of RanGTP-activated nucleosome assembly of H2A-H2B by Imp9. We use hydrogen-deuterium exchange coupled with mass spectrometry (HDX) to characterize the solution conformations and interactions of RanGTP, Imp9, and H2A-H2B, in relevant binary complexes, as well as the atypical ternary complex. We show that Imp9 stabilizes H2A-H2B beyond the direct binding site, as seen for other histone chaperones. Importantly, we reveal that RanGTP induces the release of H2A-H2B from the N-terminal repeats of Imp9, while binding at the C-terminal repeats is mostly unchanged. This partial release of H2A-H2B exposes histone- and DNA-binding sites that facilitate docking and ultimately deposition onto an assembling nucleosome. Finally, we find that RanGTP binds canonically to Imp9 in both the binary and ternary complexes but has a weaker affinity if H2A-H2B is present. A weaker affinity may ensure the release of H2A-H2B only at chromatin where RanGTP is in relatively high concentration. RanGTP thus plays an essential role in regulating the nucleosome assembly activity of H2A-H2B by Imp9.

## Results

### D-uptake of Imp9, H2A-H2B, and RanGTP alone

To probe Imp9 complexes in solution, we performed bottom-up HDX. We monitored 634 peptides covering most of the protein sequences across four time points [10, 10^2^, 10^3^, 10^4^ s] (Tables S1–2). Analysis of absolute deuterium uptake (D-uptake) of Imp9 alone correlates with the known HEAT repeat secondary structures, with most loops within and between repeats having higher D-uptake than the a- and b-helices after 10^2^ or 10^3^ s exchange (Figure S1A). Notable exceptions are the h11 and h15 loops with relatively low D-uptake, suggesting they are shielded from solvent by intra-molecular interactions. It is thus likely that Imp9 alone is adopting or at least visiting a super-helical conformation, similar to that seen in the Kap114 structure (27). The higher D-uptake of the a-helices (outer side of super-helix) compared to the b-helices (inner side of super-helix) further supports a super-helical conformation in solution. The h8, h19, and h18-h19 loops also had high D-uptake at early time points, indicating disorder. Similar analysis of H2A-H2B alone gave similar D-uptake trends to those reported for yeast histones (28). We see EX1 bimodality in many peptide spectra indicative of cooperative unfolding at the ionic strength of our experiment [200 mM NaCl, 2 mM Mg(CH_3_COO)_2_•4H_2_O]. We did not recover peptides from the N-terminal tail of H2A or H2B. Finally, for RanGTP alone, we see a lower D-uptake in the core compared to surface elements, confirming a folded, globular structure (Figure S1B). Switch 1 and 2 had high D-uptake at early time points showing flexibility. We can induce bimodality in many RanGTP peptide spectra by adding EDTA to compete away the structural magnesium; hence, the fact that bimodality was not seen in our experimental conditions suggests the RanGTP is fully saturated with the magnesium essential for GTP binding (29).

### Changes in Imp9 upon forming binary complexes with H2A-H2B or RanGTP

To characterize the interactions between Imp9 and H2A-H2B or RanGTP, we looked at the difference in D-uptake between Imp9 alone and in binary complexes, Imp9•H2A-H2B or Imp9•RanGTP, respectively. Mapping these differences onto structures of Imp9•H2A-H2B (6) and Imp9•RanGTP (manuscript co-submitted, Jiou et al. 2023) identifies known interfaces (Figure 1). Structures in Figure 1 are colored according to the difference in D-uptake summed across all time points and averaged between overlapping peptides with no significance filter. More detailed Imp9 peptide-level data are shown in subsequent figures with a two-factor significance threshold of ≥5% difference in D-uptake and a *p*-value ≤0.01 in Welch’s t-test [n=3] (Figure 2 shows N-terminal HEAT repeats h1-h7; Figure S2 shows middle HEAT repeats h8-h15; and Figure 3 shows C-terminal HEAT repeats h15-h20) (30). From these data, we see that adding H2A-H2B to Imp9 caused reduced D-uptake within the Imp9 N-terminal repeats at the h4-h5 loop, h5, and, to a lesser extent, h6 (Figure 1A, 2A). Reduced D-uptake was observed within the C-terminal repeats at the h18 loop, the start and end of the h18-h19 loop, and the C-terminal end of the h19 a-helix (Figure 1A, 3A). These regions correspond to the binding sites of H2A-H2B in Imp9, as seen in the crystal structure where a highly-constrained Imp9 super-helix wraps around H2A-H2B (6). We also see reduced D-uptake at the beginning of the disordered h19 loop (namely, residues 932-941), most of which is not visible in the structure (Figure 3A). This loop extends from the h19a helix near the DNA-binding L1 [between the first and second helices of the histone core] and L2 [between the second and third helices of the histone core] loops of H2A and H2B, respectively. It forms transient interactions with H2A-H2B that are detectable by HDX but not crystallography.

**Figure 1.**
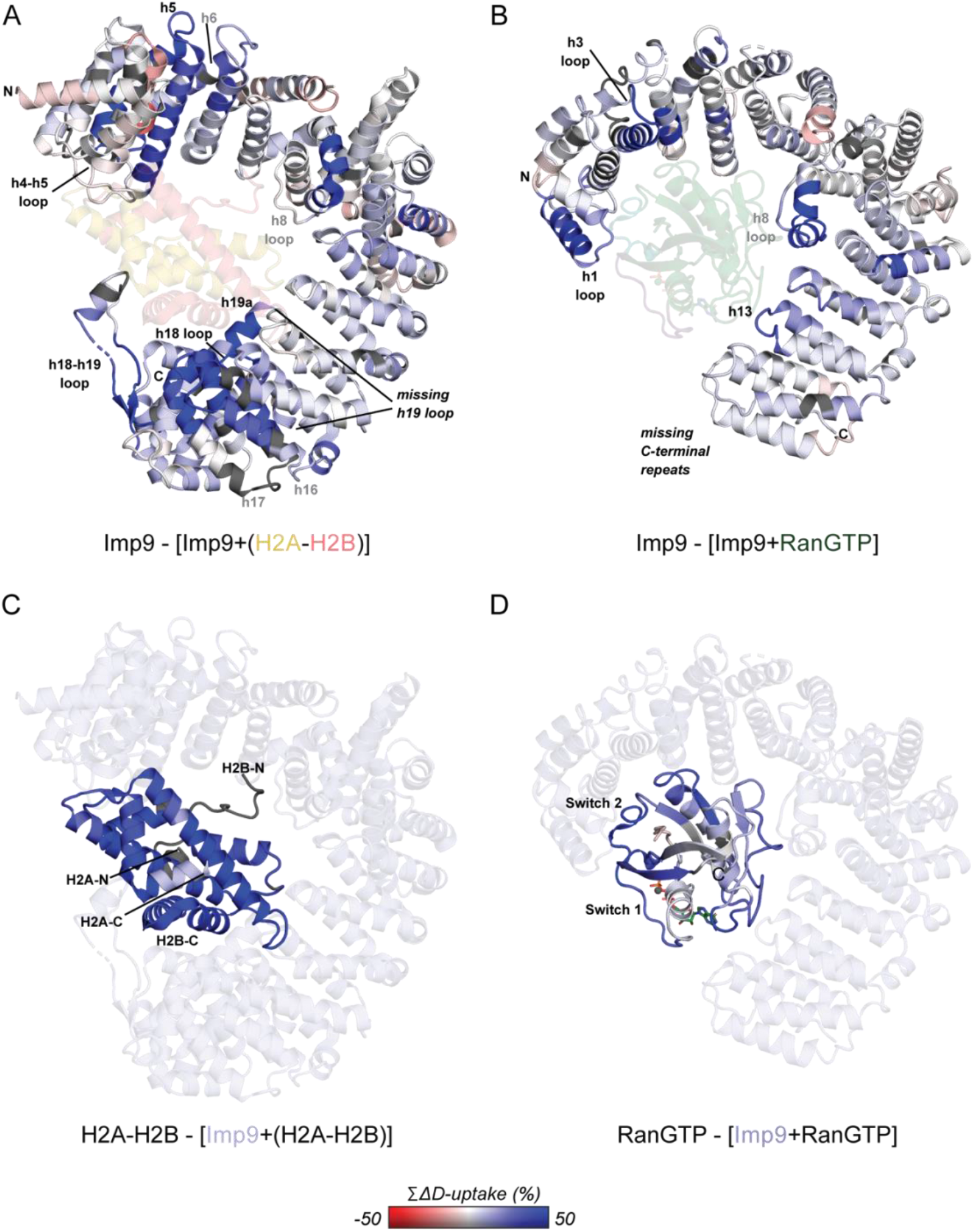
Hydrogen-deuterium exchange of binary Imp9 complexes. (**A, B**) Imp9 colored according to the change in D-uptake upon adding H2A-H2B (**A**) or RanGTP (**B**) summed across all time points. (**C, D**) H2A-H2B (**C**) or RanGTP (**D**) colored according to the change in D-uptake upon adding Imp9 summed across all time points. Imp9, H2A, H2B, and RanGTP not mapped with change in D-uptake are colored light blue, yellow, pink, or green, respectively. The scale is −50 to 50% (red to blue) and is based on DynamX residue-level scripts without statistical filtering. Residues without coverage are dark gray.

The addition of RanGTP similarly caused reduced D-uptake of Imp9. This occurred in Imp9 N-terminal repeats at h1 and the h3 loop, as well as within the middle repeats at the h8 loop and h13 repeat (Figure 1B, 2, S2). The interface between RanGTP and importins is extremely conserved among paralogs and orthologs, and typically involves these structural elements (17). The structure of Imp9•RanGTP confirms the relevance of these elements specifically for Imp9 (manuscript co-submitted, Jiou et al. 2023). The structure does not resolve repeats 16-20 and we see no difference in D-uptake for these repeats (Figure 3A), clearly showing that these C-terminal repeats are not involved in binding RanGTP. Beyond the known interfaces, we also see reduced D-uptake with either H2A-H2B or RanGTP in the Imp9 h8 loop, h16-h17 repeats, and the flexible/disordered h19 loop (Figure 1A, B, 2A, 3A). The larger reduction in D-uptake for h16-h17 with H2A-H2B than RanGTP suggests it is a hinge that positions the C-terminal repeats for histone-binding, reinforcing the additional constraint on the C-terminal repeats in Imp9 bound by H2A-H2B compared to RanGTP.

**Figure 2.**
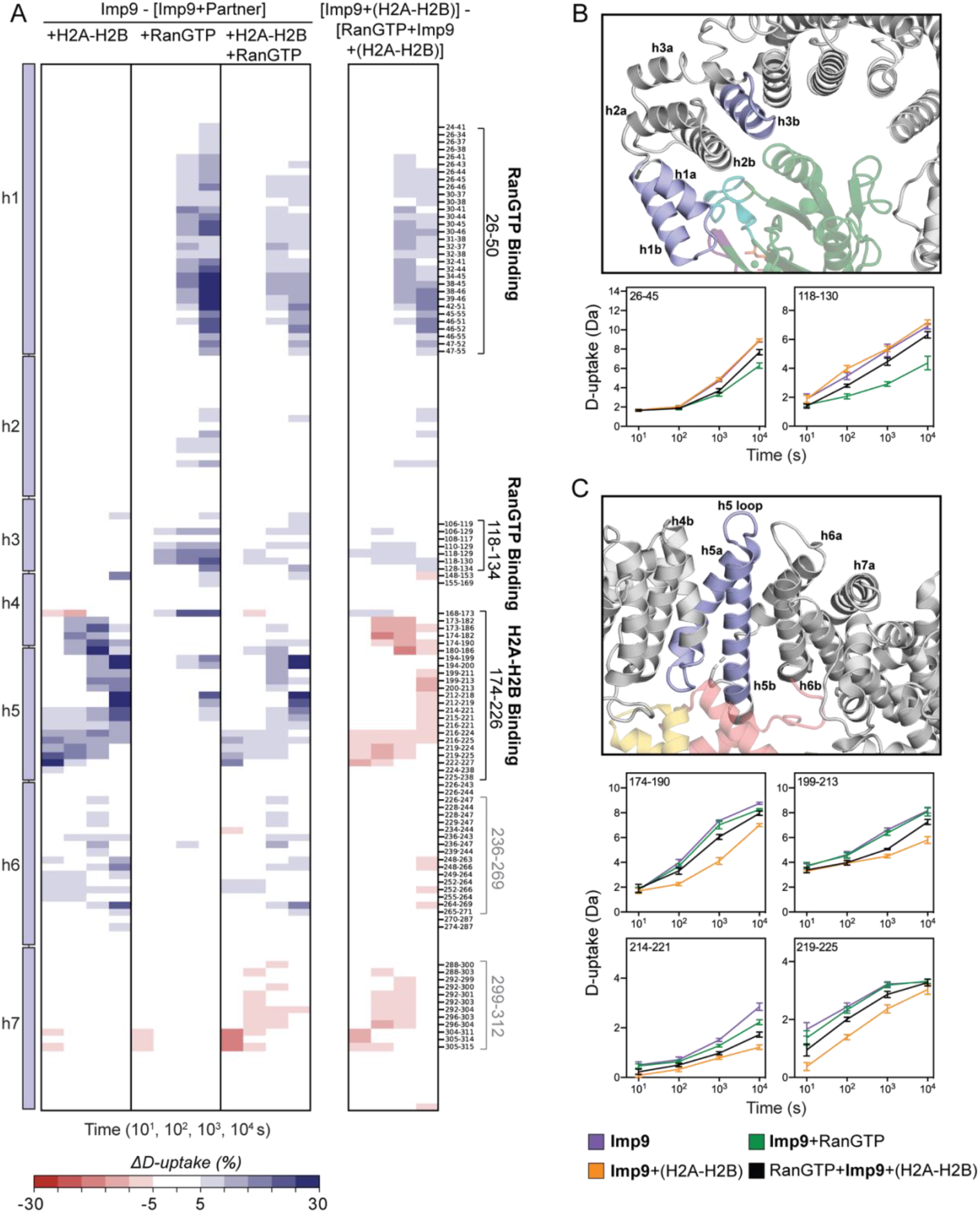
D-uptake differences in the N-terminal HEAT repeats of Imp9. (**A**) Differences in D-uptake between Imp9 alone and Imp9+(H2A-H2B), Imp9+RanGTP, or RanGTP+Imp9+(H2A-H2B), in **Panels 1,2**, and **3**, respectively. Difference in D-uptake between Imp9+(H2A-H2B) and RanGTP+Imp9+(H2A-H2B) is shown in **Panel 4**. Residue boundaries and peptides from binding sites (black) and regions of conformational change (gray) are noted. Blue/Red coloring indicates a difference ≥5% with a *p*-value ≤0.01 in Welch’s t-test [n=3]. (**B**) Interface between the N-terminal HEAT repeats of Imp9 (light blue) and RanGTP (green) with example D-uptake plots of peptides from Imp9 residues 26-50 and 118-134 (colored light purple on structure). (**C**) Interface between the N-terminal HEAT repeats of Imp9 (light blue) and H2A-H2B (H2A in yellow, H2B in pink) with example D-uptake plots of peptides from Imp9 residues 174-226 (colored light purple on structure). For B and C, D-uptake plots show Imp9 alone (purple), Imp9+(H2A-H2B) (orange), Imp9+RanGTP (green), and RanGTP+Imp9+(H2A-H2B) (black). Error bars are ±2SD with n=3. The y-axis is 80% of the maximum theoretical D-uptake, assuming the complete back exchange of the N-terminal residue.

### Changes in H2A-H2B or RanGTP upon forming binary complexes with Imp9

We can further compare the D-uptake of peptides from H2A-H2B or RanGTP alone to those in the binary Imp9 complexes (Figure 1C, D). H2A-H2B and RanGTP peptide-level data with statistical filters are shown in subsequent figures (Figure S3 shows H2A-H2B, and Figure 4 shows RanGTP). For H2A-H2B, adding Imp9 caused reduced D-uptake throughout the entire folded regions of the heterodimer (Figure S3). These include the histone-fold domain composed of α1, α2, and α3 helices from both H2A and H2B, as well as the H2B C-terminal helix. The only region we see no change in D-uptake is the C-terminal tail of H2A (Figure S3A). This tail is maximally deuterated in H2A-H2B alone and Imp9•H2A-H2B, suggesting it is disordered in both conditions. The somewhat global stabilization of H2A-H2B by Imp9 is like that seen with histone chaperone Nap1 or an increase in ionic strength (28). In all three cases, the reduction in D-uptake in H2A is less than in H2B, and the reduction is most pronounced in the H2A α2 and the H2B α2, α3, and C-terminal helices (Figure S3). Given that Imp9 and Nap1 have distinct modes of interaction with H2A-H2B (6, 28), this shared pattern cannot be attributed to direct binding interactions alone. Rather, it must involve shared allosteric effects that stabilize hydrogen bonds outside of the binding interfaces. These allosteric effects likely occur as Imp9, Nap1, or salt, reduce the cooperative unfolding of the H2A-H2B heterodimer (28). Cooperative unfolding is indicated by EX1 bimodality in the H2A-H2B peptide spectra, and this bimodality disappears or is delayed when Imp9, Nap1, or salt, is added (28). Figure S4 shows an example peptide from H2B with EX1 bimodality in H2A-H2B alone that disappears or is delayed when Imp9 is present. The peptide is from a region of H2B that does not directly contact Imp9.

The identification of Imp9-binding regions of RanGTP is more straightforward. Adding Imp9 caused reduced D-uptake primarily at Switch 2, suggesting it to be the primary RanGTP region for contacting Imp9 (Figure 4A). Reduced D-uptake was also seen for RanGTP Switch 1 and, to a lesser extent, surface loops composed of residues 93-120 and 121-158. In the many homologous importin-bound RanGTP structures, these regions of RanGTP interface with the importin protein (17). The structure of Imp9•RanGTP specifically corroborates their relevance for Imp9 (manuscript co-submitted, Jiou et al. 2023). More specifically, the RanGTP Switch 1 contacts Imp9 h1 and h15; Switch 2 contacts Imp9 h1 and h2; residues 93-120 contacts Imp9 h3; and residues 121-158 contacts the loops of Imp9 h8 and h12-h14 (Figure 1D). The solution and structural data are thus consistent in identifying the key contacts in binary Imp9 complexes.

### RanGTP reduces occupancy of H2A-H2B at the Imp9 N-terminal HEAT repeats

Concurrent with the binary Imp9 complexes, we also performed HDX on the ternary complex containing RanGTP, Imp9, and H2A-H2B. We have previously shown RanGTP•Imp9•H2A-H2B to be a stable 1:1:1 complex that, unlike Imp9•H2A-H2B, has nucleosome assembly activity (6). The question was: how does the binding of RanGTP allow H2A-H2B to be released to a nucleosome intermediate? To answer this, we examined the difference in D-uptake between the binary and ternary complexes from the perspective of Imp9 (Figure 2, 3, S2), H2A-H2B (Figure S3), and RanGTP (Figure 4). We applied statistical thresholds as above and show regions of interest on binary structures as the ternary Imp9 structure has not been solved.

**Figure 3.**
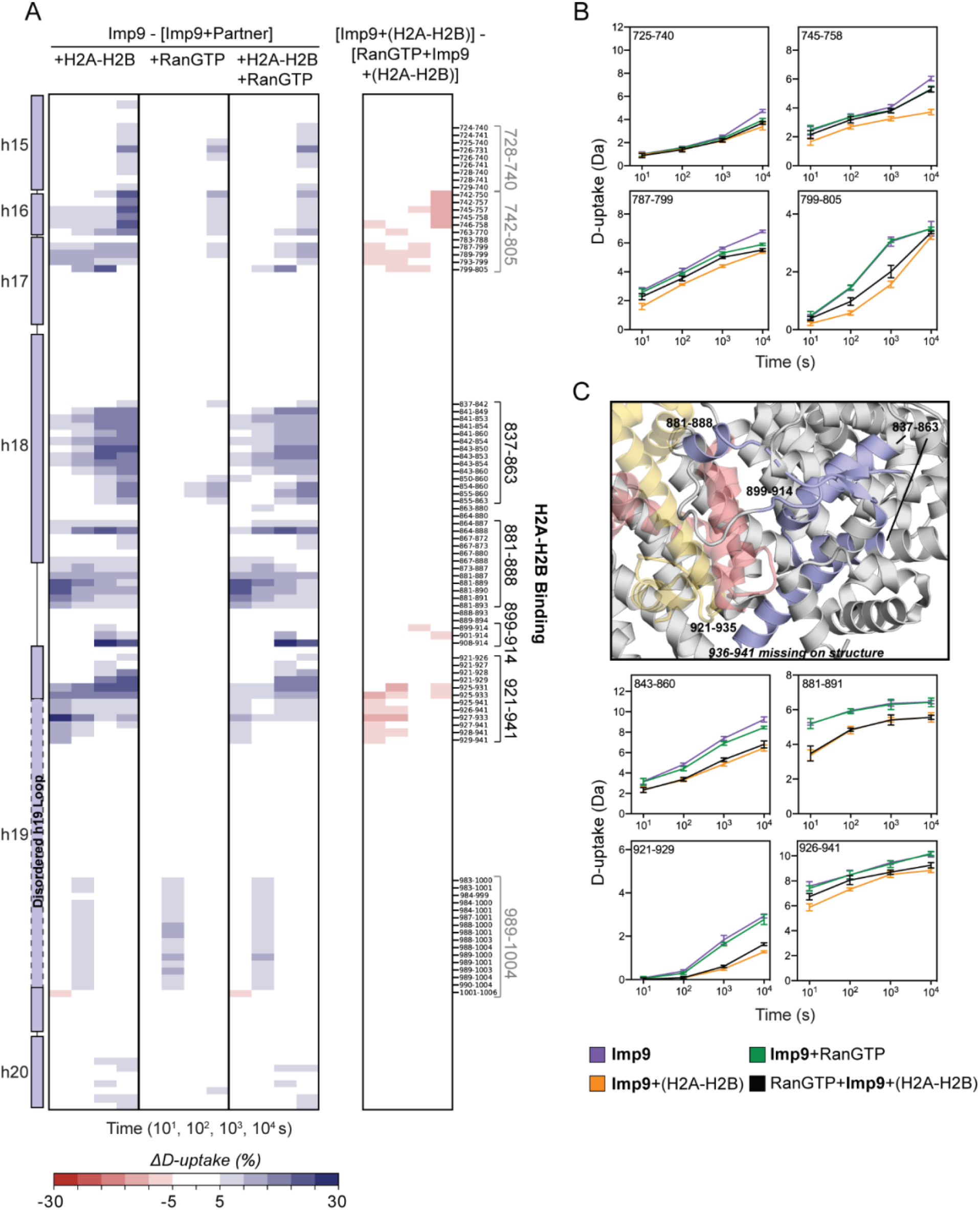
D-uptake differences in the C-terminal HEAT repeats of Imp9. (**A**) Difference in D-uptake between Imp9 and Imp9+(H2A-H2B), Imp9+RanGTP, or RanGTP+Imp9+(H2A-H2B) in **Panel 1**, **2**, and **3**, respectively. Difference in D-uptake between Imp9+(H2A-H2B) and RanGTP+Imp9+(H2A-H2B) is shown in **Panel 4**. Residue boundaries and peptides from binding sites (black) and regions of conformational change (gray) are noted. Blue/Red coloring indicates a difference ≥5% with a *p*-value ≤0.01 in Welch’s t-test [n=3]. (**B**) Region of conformational change in the C-terminal HEAT repeats of Imp9 upon binding H2A-H2B and/or RanGTP with example D-uptake plots for peptides from Imp9 residues 728-805. (**C**) Interface between the C-terminal HEAT repeats of Imp9 (light blue) and H2A-H2B (H2A in yellow, H2B in pink) with example D-uptake plots of peptides from Imp9 residues 837-863, 881-888, 899-914, and 921-941 (colored light blue on structure). For B and C, D-uptake plots show Imp9 alone (purple), Imp9+(H2A-H2B) (orange), Imp9+RanGTP (green), and RanGTP+Imp9+(H2A-H2B) (black). Error bars are ±2SD with n=3. The y-axis is 80% of the maximum theoretical D-uptake, assuming the complete back exchange of the N-terminal residue.

Comparison of Imp9•H2A-H2B to RanGTP•Imp9•H2A-H2B revealed that RanGTP alters H2A-H2B binding within the N-terminal repeats of Imp9, while binding within the C-terminal repeats is relatively unchanged. Adding RanGTP caused an increase in D-uptake for h5 and the preceding h4-h5 loop, indicating reduced H2A-H2B occupancy at this N-terminal site (Figure 2A). This was observed for 18 of the 20 peptides spanning Imp9 residues 174 to 226, with the two exceptions being short peptides (residues 194-199 and 194-200) that cover the h5 loop that is situated on the outer side of the Imp9 super-helix. This region and D-uptake plots of four representative peptides are highlighted in Figure 2C. In contrast, adding RanGTP did not change the D-uptake of the h18 loop or the start of the h18-h19 loop (residues 881-888), indicating unchanged occupancy of H2A-H2B at these major C-terminal contacts (Figure 3A). We do, however, observe increased D-uptake of a nearby region, including the end of the h18-h19 loop (residues 899-914), the end of the h19 a-helix, and the start of the h19 loop. This increase suggests H2A-H2B contacts with Imp9 in this region are weakened or altered by the addition of RanGTP (Figure 3A). Example D-uptake plots are shown in Figure 3C. Beyond the Imp9 H2A-H2B-binding sites, RanGTP also caused increased D-uptake at the h16-h17 hinge (Figure 3A). D-uptake plots very clearly show that in the ternary complex this h16-h17 hinge exchanges like Imp9•RanGTP rather than Imp9•H2A-H2B (Figure 3B). The RanGTP-induced loss of H2A-H2B binding at the N-terminal repeats of Imp9 seems to cause an opening of the h16-h17 hinge such that the C-terminal repeats of Imp9, where H2A-H2B remains bound, are less constrained. In the ternary Imp9 complex, the N-terminal repeats resemble the RanGTP-bound binary complex, while the C-terminal repeats resemble the H2A-H2B-bound binary complex.

The RanGTP-induced release of H2A-H2B from the N-terminal repeats of Imp9 makes H2A-H2B more accessible to solvent and/or for nucleosome assembly. The ‘end’ of H2A-H2B released contains the L2 loop of H2A and the L1 loop of H2B that binds DNA in the nucleosome (1). It is also close to the H2A C-terminal region that drives the docking of H2A-H2B onto (H3-H4)2 in the nucleosomal histone octamer (1). A reasonable prediction is that this end of H2A-H2B would show an increase in D-uptake when RanGTP was added to Imp9•H2A-H2B, concomitant with the release from Imp9. This increase, however, was not observed, and there was little to no change in H2A-H2B D-uptake between the binary and ternary complexes (Figure S3). The likely reason for this is the allosteric stabilization of H2A-H2B by Imp9, with the interaction between a single end of H2A-H2B and the Imp9 C-terminal repeats being sufficient for the full allosteric effect. This is precedented by Nap1, as it elicits the same allosteric effect and binds a single end of H2A-H2B (28). The expected changes in H2A-H2B are likely below our stringent statistical thresholds as the changes are small compared to the allosteric effect, and because not all H2A-H2B is released from Imp9 (see next paragraph). Notably, two very small changes in D-uptake were detected. The first was a decrease in the D-uptake of H2A and H2B α2 helices near the released end of H2A-H2B (Figure S3B). This may be due to new (but weak) interactions between released H2A-H2B and Imp9 or RanGTP, similar to those seen in the homologous ternary complex structure of RanGTP•Kap114•H2A-H2B (manuscript cosubmitted, Jiou et al. 2023). The second was an increase in D-uptake of H2A α1 (Figure S3A), potentially due to altered interactions with the nearby start of the h19 loop for which we also detect changes in the ternary complex (Figure 3A).

### H2A-H2B compromised the affinity between RanGTP and Imp9

We can also consider the effect that H2A-H2B has on RanGTP binding by looking at the RanGTP-binding site in Imp9 (Figure 2A). This revealed that the presence of H2A-H2B reduces the occupancy of RanGTP on Imp9. This is most clearly indicated by the reduction in D-uptake upon adding RanGTP being greater for the binary complex than the ternary (Figure 2A, compare panels 2 and 3). This was seen for the RanGTP-binding site in Imp9 (h1 and h3) and is evident in example D-uptake plots (Figure 2A, B; compare black and green traces). This trend was also observed for RanGTP peptides involved in binding Imp9. More change in the binary complex than the ternary occurred at Switch 1, Switch 2, and residues 93-120 and 121-158 (Figure 4A, B). Importantly, it is only the occupancy of RanGTP that is altered and not the actual contacts as the pattern of D-uptake remained the same between the binary and ternary complexes for both Imp9 and RanGTP peptides (Figure 2A, 4A). Using Imp9 peptides from the RanGTP-binding site, we calculated the change in D-uptake of the ternary complex to be ~55% of the binary complex (Table S3). As RanGTP was limiting in our experiment and thus 100% bound in the binary complex, it must be only 55% bound in the ternary. This predicts an affinity of ~200 nM, which is weaker than the reported single-digit nanomolar affinities for several importin and RanGTP pairs (31, 32). To test the prediction, we used fluorescence polarization to measure the affinity of RanGTP for Imp9•H2A-H2B and Kap114•H2A-H2B (Figure S5). The measured affinities were close to the predicted value at 150 and 270 nM. Notably, these are at least 100-fold weaker than the affinity of RanGTP for only Imp9 or Kap114 measured using the same approach (manuscript co-submitted, Jiou et al. 2023). These data clearly show that although binding remains tight, the presence of H2A-H2B does compromise the affinity of RanGTP for Imp9, consistent with the HDX result. A weaker affinity for Imp9/Kap114•H2A-H2B than Imp9/Kap114 alone may ensure that H2A-H2B is only released from the importin near chromatin where the RanGTP concentration is higher (20, 21).

**Figure 4.**
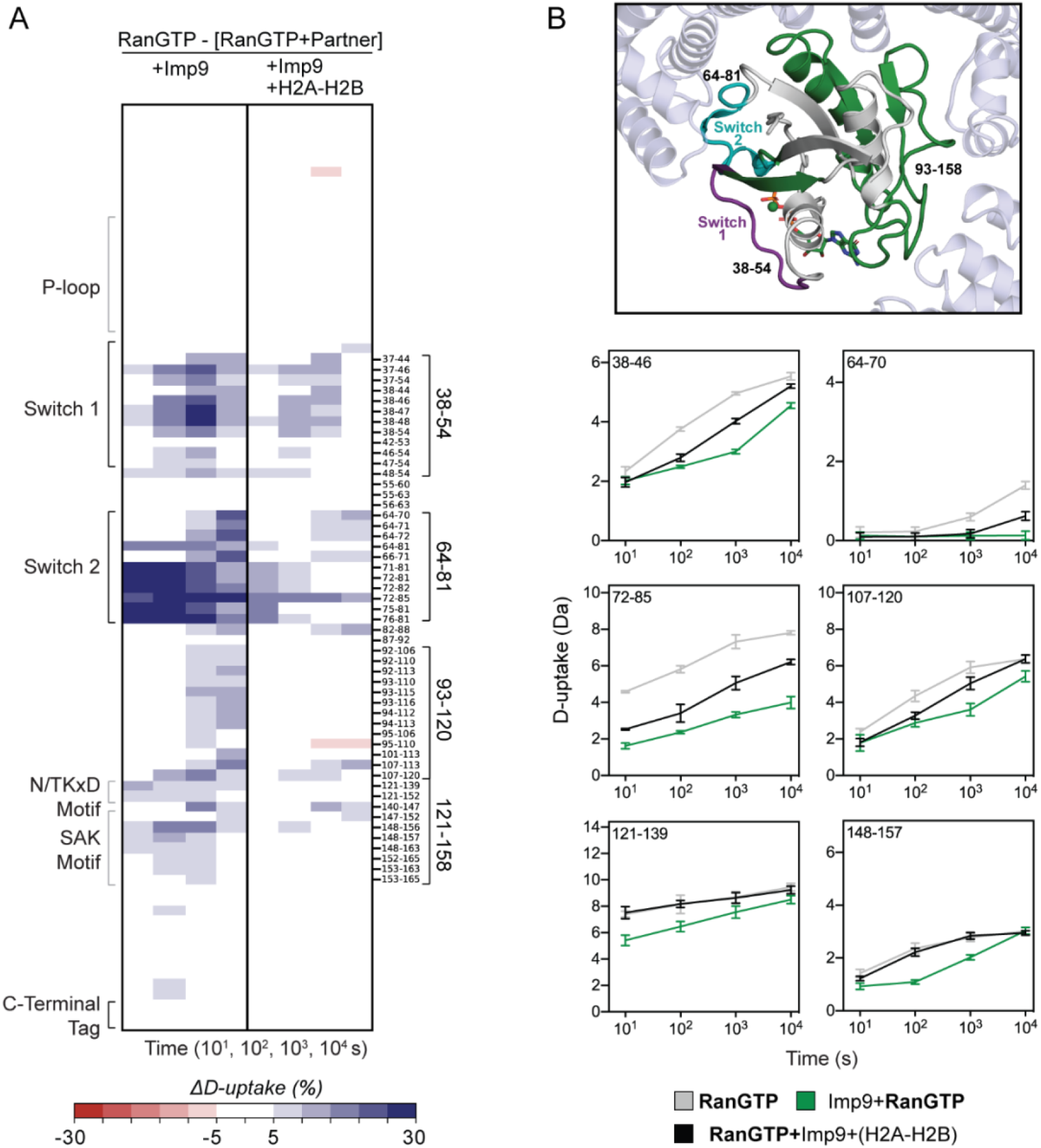
D-uptake differences in RanGTP. (**A**) Differences in D-uptake between RanGTP alone and Imp9+RanGTP or RanGTP+Imp9+(H2A-H2B) in **Panels 1**, **2**, respectively. Residue boundaries and peptides from binding sites (black) are noted. Blue/Red coloring indicates a difference ≥5% with a *p*-value ≤0.01 in Welch’s t-test [n=3]. (**B**) Interface between RanGTP (green) and Imp9 (light blue) with example D-uptake plots for peptides from RanGTP residues 38-54, 64-81, 93-120, and 121-158 (colored green on structure). D-uptake plots show RanGTP alone (gray), Imp9+RanGTP (green), and RanGTP+Imp9+(H2A-H2B) (black). Error bars are ±2SD with n=3. The y-axis is 80% of the maximum theoretical D-uptake, assuming the complete back exchange of the N-terminal residue.

## Discussion

We have identified the physical and thermodynamic differences between the binary Imp9•H2A-H2B and Imp9•RanGTP complexes, and the non-canonical RanGTP•Imp9•H2A-H2B ternary complex. The binding of RanGTP releases H2A-H2B from the N-terminal repeats of Imp9 with little to no effect on the binding at the C-terminal repeats. The variable conformation of the hinge between repeats 16 and 17 means the N-terminal repeats of Imp9 resemble Imp9•RanGTP, while the C-terminal repeats resemble Imp9•H2A-H2B. Structural studies with yeast Kap114 are similar and suggest this mechanism is conserved across species (manuscript co-submitted, Jiou et al. 2023). When both the N-terminal and C-terminal repeats of Imp9/Kap114 are bound to H2A-H2B, Imp9/Kap114 acts as a histone chaperone that shields H2A-H2B and prevents interaction with DNA or an assembling nucleosome. When only the C-terminal repeats of Imp9/Kap114 are bound to H2A-H2B due to the co-binding of RanGTP, Imp9 switches to a nucleosome assembly factor that deposits H2A-H2B in a nucleosome. The RanGTP-induced exposure of new H2A-H2B surfaces facilitates its interaction with DNA and/or other histones. While we have tested H2A-H2B release to an assembling nucleosome, it is also feasible that H2A-H2B is transported between Imp9•RanGTP and another histone chaperone, such as Nap1 or FACT, or chromatin remodeling factor in the nucleus (28, 33). Kap114 binds Nap1 directly (34, 35), and Imp9 co-elutes with Nap1L1 and Nap1L4 in mammalian cells, although a direct interaction has not been probed (36–39).

We have shown that the affinity between RanGTP and Imp9/Kap114 is weaker in the presence of H2A-H2B, even though the molecular determinants of the interaction are essentially identical. A weaker affinity implies a requirement for higher RanGTP concentrations if RanGTP is to bind Imp9/Kap114 occupied by H2A-H2B, as opposed to other cargo with canonical release mechanisms (22, 40). These higher concentrations are likely near chromatin where the Ran guanine nucleotide exchange factor RCC1 resides (20, 21), ensuring that RanGTP causes the partial release of H2A-H2B right where it is needed (20, 38, 39). It is unclear precisely how RanGTP binds when it first encounters Imp9/Kap114•H2A-H2B. The Imp9 residues that bind RanGTP are accessible in the binary complex and could possibly engage the GTPase weakly in the initial encounter when H2A-H2B is fully engaged. Imp9/Kap114 may need to sample a super-helical conformation that truly results in stable RanGTP binding. Alternatively, RanGTP may only bind to a transient conformation of Imp9/Kap114•H2A-H2B where the N-terminal repeats of Imp9 detach from H2A-H2B (41). The necessity for conformational change in Imp9/Kap114•H2A-H2B to bind RanGTP is consistent with the weakened affinity for RanGTP binding to Imp9/Kap114•H2A-H2B compared to Imp9/Kap114 alone.

Our observation that Imp9 causes stabilization of H2A-H2B beyond the direct binding interface is remarkably like studies with other H2A-H2B chaperones, namely Nap1 and FACT (28, 33). The degree of stabilization in different H2A-H2B secondary structures is similar in all cases, despite the chaperones recognizing the histones in unique ways. The generality of this effect is reinforced by increases in ionic strength causing the same patterns of stabilization (28). The cooperative unfolding of yeast H2A-H2B has been reported (28), and we now see the same behavior for frog H2A-H2B. Chaperone binding reduces this unfolding and accounts for the stabilization or allosteric effect observed. Unfolding of H2A-H2B in its free state may partially account for the requirement for chaperones. H3-H4 chaperones DAXX and Caf1 act similarly, suggesting this phenomenon also applies to H3-H4 (42, 43). Another shared feature between Imp9/Kap114 and histone chaperones is the presence of an acidic stretch of amino acids. FACT sets a precedent that these acidic stretches act as a DNA mimic during the transfer of H2A-H2B to a nucleosome (33). The acidic h19 loop of Imp9/Kap114 may play a similar role, especially as it is located right near DNA-binding loops of H2A-H2B.

The combination of HDX and cryo-EM is a valuable approach to studying many protein systems (33, 44–46). HDX provides solution relevance and thermodynamic insights to the high-resolution structures produced by cryo-EM. The Imp9•RanGTP structure (manuscript co-submitted, Jiou et al. 2023) corroborates the binding interfaces identified by HDX, while HDX informs on the weakened affinity in the presence of H2A-H2B. The mechanism of RanGTP-induced nucleosome assembly activity of Imp9 derived from HDX is in line with that for Kap114 based on structures (manuscript co-submitted, Jiou et al. 2023). The conserved mechanism involves Imp9/Kap114 chaperoning H2A-H2B from the cytoplasm, through the nuclear pore complex, and into the nucleus, where RanGTP binding can facilitate H2A-H2B assembly into chromatin. Future work will place these import and deposition activities more clearly in the context of other chromatin assembly and disassembly pathways in the cell.

## Materials and Methods

### Constructs, Protein Expression and Purification

Human Importin-9 and yeast RanGTP (*Saccharomyces cerevisiae* Gsp1 residues 1-179 with Q71L) were expressed and purified as in Padavannil et al, 2019 (6). Kap114 was expressed and purified as in Jiou et al, 2023 (co-submitted manuscript). Histones were obtained from The Histone Source and refolded as described in Luger et al, 1997 (1).

### Hydrogen-Deuterium Exchange Mass Spectrometry

Samples were prepared from 25 μM stocks of Imp9, frog H2A-H2B, RanGTP, Imp9•H2A-H2B (1:1), Imp9•RanGTP (1:0.8), and RanGTP•Imp9•H2A-H2B (0.8:1:1) in 20 mM HEPES pH 7.5, 200 mM NaCl, 2 mM Mg(CH_3_COO)_2_•4H_2_O, 2 mM TCEP, 10% (v/v) glycerol, pH 7.5. These stocks were equilibrated at 25°C for 20 min. Control samples were diluted 1:24 with this buffer containing H_2_O. For exchange samples, the same buffer containing D_2_O was used for 1:24 dilution (pH_r_ead = 7.1 at 25°C; final D_2_O of 96%). Exchange proceeded at 25°C for 10, 10^2^, 10^3^, or 10^4^ s. Exchange was quenched by mixing samples 1:1 with cooled 0.8% (v/v) formic acid, 2 M urea, pH 1.7 (1:1 mix had a final pH of 2.3 at 0°C) and flash frozen in liquid nitrogen. Samples were prepared in triplicate and stored at −80°C.

Samples were thawed for 50 s immediately before injection into a Waters™ HDX manager in line with a SYNAPT G2-Si. In the HDX manager, samples were digested by *Sus scrofa* Pepsin A (Waters™ Enzymate BEH) at 15°C and the peptides trapped on a C4 pre-column (Waters™ Acquity UPLC Protein BEH C4) at 1°C using a flowrate of 100 μl/min for 3 min. The chromatography buffer was 0.1% (v/v) formic acid. Peptides were separated over a C18 column (Waters™ Acquity UPLC BEH) at 1°C and eluted with a linear 3-40% (v/v) acetonitrile gradient using a flowrate of 40 μl/min for 7 min. Samples were injected in a random order.

Mass spectrometry data were acquired using positive ion mode in either HDMS or HDMS^E^ mode. Peptide identification of water-only control samples was performed using data-independent acquisition in HDMS^E^ mode. Peptide precursor and fragment data were collected via collision-induced dissociation at low (6 V) and high (ramping 22-44 V) energy. HDMS mode was used to collect low energy ion data for all deuterated samples. All samples were acquired in resolution mode. Capillary voltage was set to 2.4 kV for the sample sprayer. Desolvation gas was set to 650 L/h at 175°C. The source temperature was set to 80°C. Cone and nebulizer gas was flowed at 90 L/h and 6.5 bar, respectively. The sampling cone and source offset were both set to 30 V. Data were acquired at a scan time of 0.4 s with a range of 100–2000 m/z. Mass correction was done using [Glu1]-fibrinopeptide B as a reference. For ion mobility, the wave velocity was 700 ms^-1^ and the wave height was 40 V.

Raw data of Imp9, H2A-H2B, and RanGTP water-only controls were processed by PLGS (Waters™ Protein Lynx Global Server 3.0.3) using a database containing *S. scrofa* Pepsin A, RanGTP, H2A-H2B, and Imp9. In PLGS, the minimum fragment ion matches per peptide was 3 and methionine oxidation was allowed. The low and elevated energy thresholds were 250 and 50 counts, respectively, and overall intensity threshold was 750 counts. DynamX 3.0 was used to search the deuterated samples for peptides with 0.3 products per amino acid and 1 consecutive product found in 2 out of 4-5 controls. Data were manually curated. Structural images were made using scripts from DynamX in PyMOL version 2.5 (Schrödinger, http://www.pymol.org/pymol); and heatmaps were made using in-house Python scripts.

To allow access to the HDX data from this study, the HDX data summary table (Table S1) and the HDX state data table (Table S2) are included (47). Theoretical maximum D-uptake used in percent calculations was determined as follows: 0.96 × (residues in peptide – 1 for N-terminal residue – the number of prolines not at the N-terminal position). Back exchange was calculated using peptides from the Imp9, H2A-H2B, or RanGTP only samples that had plateaued (<2% difference in D-uptake at 10^3^ and 10^4^ s) and had a D-uptake >40%. The number reported in Table S1 is (100 – the average % D-uptake for these peptides at 10^4^ s). The number of peptides used in the calculation was 83, 16, 6, and 9 for Imp9, H2A, H2B, and RanGTP, respectively. The raw mass spectrometry data are available from ProteomeXchange via the PRIDE partner repository with identifier PXD037571 (48–50).

### Fluorescent Polarization

Fluorescence polarization (FP) assays were performed in a 384-well format as previously described (17, 51). Kap114, Imp9, and RanGTP were dialyzed overnight into 20 mM HEPES pH 7.5, 150 mM NaCl, 2 mM MgCl2, 2 mM TCEP, 10% (v/v) glycerol. 12 μM RanGTP was serially diluted with buffer and mixed with 80 nM ^XFD488^H2A-H2B and 60nM Imp9/Kap114 (1:1 volume with final volume of 15 μL, 2X of final concentration). Triplicate reactions were analyzed in black-bottom plates (Corning), and data were collected in a CLARIOstar plus plate reader (BMG Labtech) equipped with dichroic filter LP 504. Measurement was performed with top optics with an excitation range of 482-16 nm and emission range of 530-40 nm, 50 flashes per well at a focal height of 5.1 mm with a 0.1 s settling time. The gainwas adjusted using a well with only ^XFD488^H2A-H2B target mP of 200 and kept constant for the rest of the measurements. Data were analyzed in PALMIST (52) and fitted with a 1:1 binding model, using the error surface projection method to calculate the 95% confidence intervals of the fitted data. Fitted data were exported and plotted in GUSSI (53).

## Acknowledgments

We thank Divyasri Damacharla for initial HDX troubleshooting. This work was funded by NIGMS of NIH under Awards R35GM133751 (S.D.), R35GM141461 (Y.M.C.), R01GM069909 (Y.M.C.); the Welch Foundation Grants AT-2059-20210327 (S.D.), I-1532 (Y.M.C.); an NSF MRI 2018188 (S.D.); support from the Alfred and Mabel Gilman Chair in Molecular Pharmacology and the Eugene McDermott Scholar in Biomedical Research (Y.M.C.); and the Gilman Special Opportunities Award (H.Y.J.F.).

## Supplemental Information

**Figure S1:**
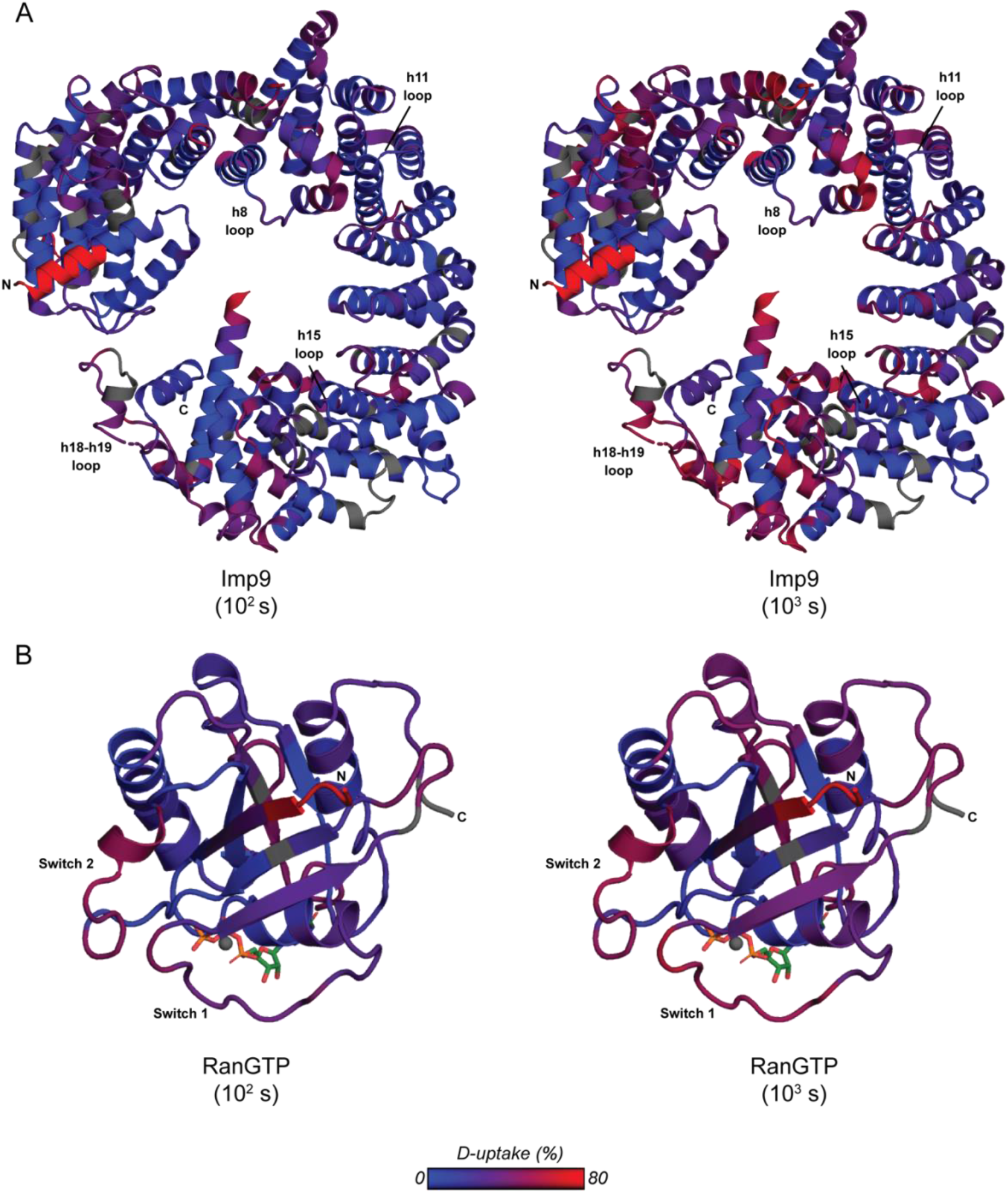
Deuterium uptake of Imp9 alone (**A**) or RanGTP alone from Kap114•RanGTP structure solved in manuscript co-submitted, Jiou et al. 2023 (**B**) after 10^2^ s (left) and 10^3^ s (right) exchange. Proteins are colored from blue to red with a 0 to 80% D-uptake scale. Coloring was done using DynamX residue-level scripts without statistical filtering. Residues without coverage are dark gray.

**Figure S2:**
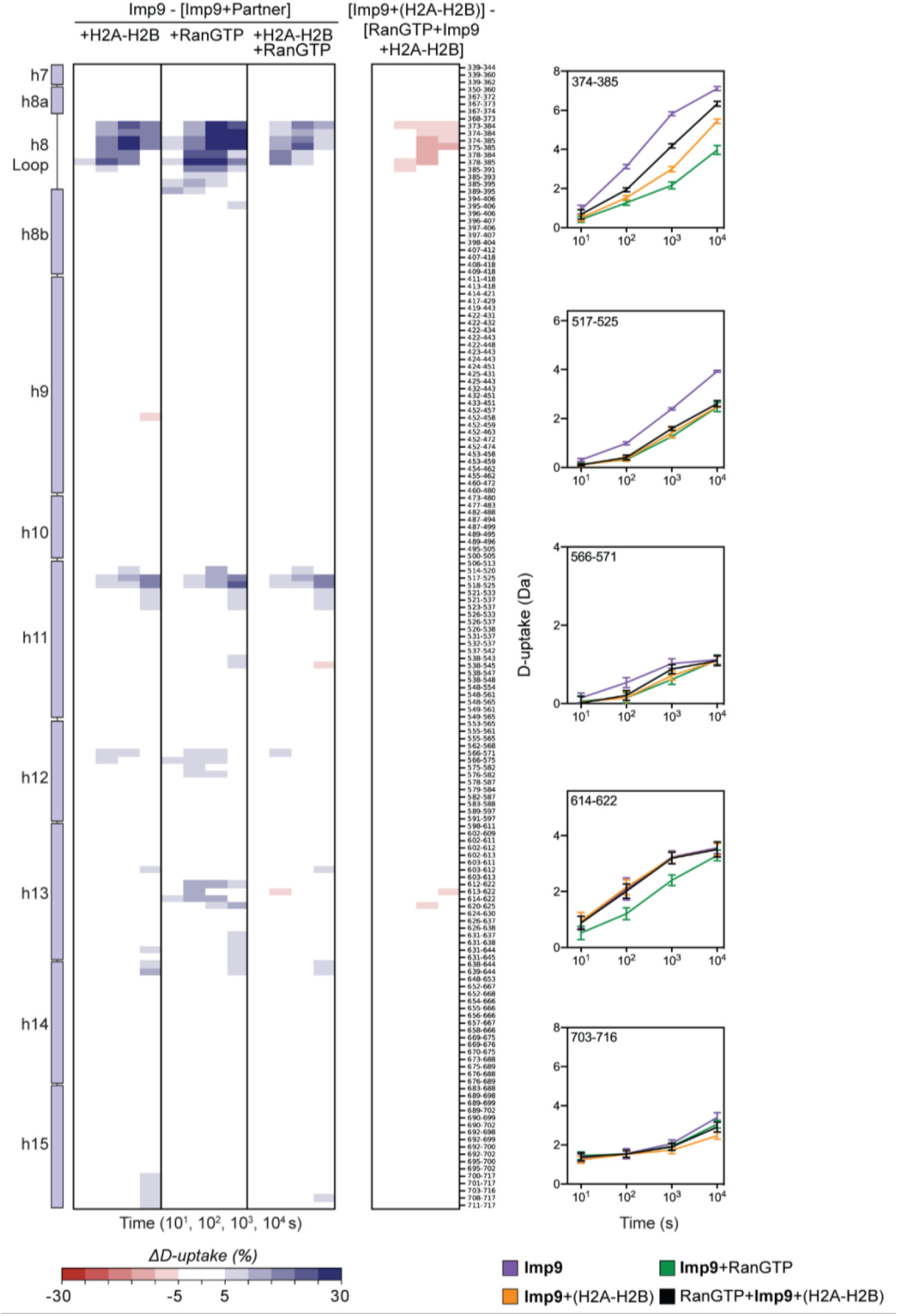
D-uptake differences in the middle HEAT repeats of Imp9. Differences in D-uptake between Imp9 alone and Imp9+(H2A-H2B), Imp9+RanGTP, or RanGTP+Imp9+(H2A-H2B) in **Panels 1**, **2**, and **3**, respectively. Difference in D-uptake between Imp9+(H2A-H2B) and RanGTP+Imp9+(H2A-H2B) is shown in **Panel 4**. Blue/Red coloring indicates a difference ≥5% with a *p*-value ≤0.01 in Welch’s t-test [n=3]. D-uptake plots of Imp9 peptides show Imp9 alone (purple), Imp9+(H2A-H2B) (orange), Imp9+RanGTP (green), and RanGTP+Imp9+(H2A-H2B) (black). Error bars are ±2SD with n=3. The y-axis is 80% of the maximum theoretical D-uptake, assuming the complete back exchange of the N-terminal residue.

**Figure S3:**
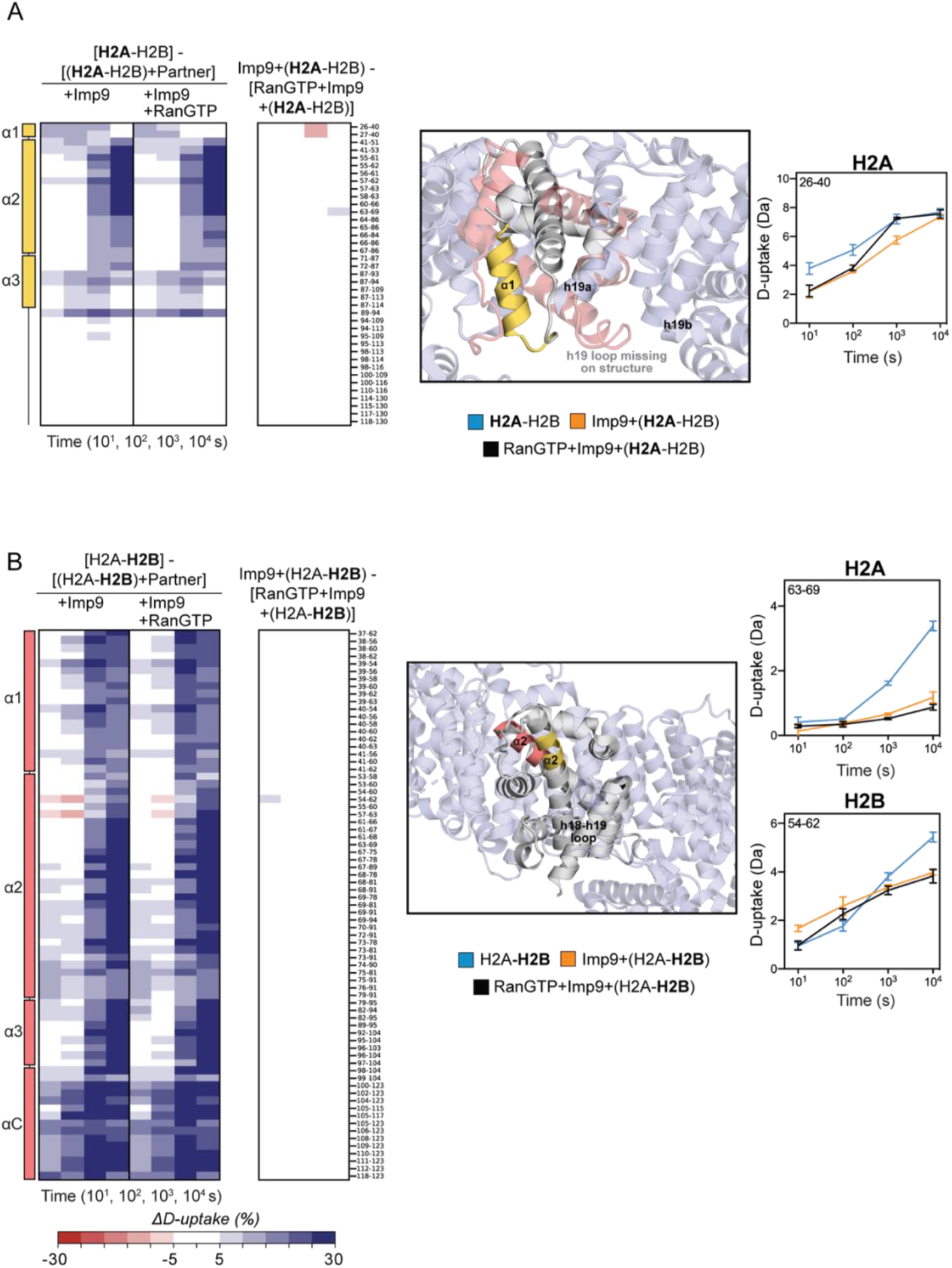
D-uptake differences in H2A (colored yellow in structure) and H2B (colored pink in structure). (**A**) Differences in D-uptake between H2A alone and Imp9+(H2A-H2B) or RanGTP+Imp9+(H2A-H2B) in **Panel 1**, and **2**, respectively. Difference between Imp9+(H2A-H2B) and RanGTP+Imp9+(H2A-H2B) shown in **Panel 3**. Zoom of H2A α1 in the Imp9•H2A-H2B structure with the D-uptake plot of H2A peptide, residues 26-40 (yellow). (**B**) Differences in D-uptake between H2B alone and Imp9+(H2A-H2B) or RanGTP+Imp9+(H2A-H2B) in **Panel 1**, and **2**, respectively. Difference between Imp9+(H2A-H2B) and RanGTP+Imp9+(H2A-H2B) shown in **Panel 3**. Zoom of H2A-H2B in the Imp9•H2A-H2B structure with D-uptake plots of H2A peptide, residues 63-69 (yellow) and H2B peptide, residues 54-62 (pink). Blue/Red coloring indicates a difference ≥5% with a *p*-value ≤0.01 in Welch’s t-test [n=3]. D-uptake plots show H2A-H2B alone (blue), Imp9+(H2A-H2B) (orange), and RanGTP+Imp9+(H2A-H2B) (black). Error bars are ±2SD with n=3. The y-axis is 80% of the maximum theoretical D-uptake, assuming the complete back exchange of the N-terminal residue.

**Figure S4:**
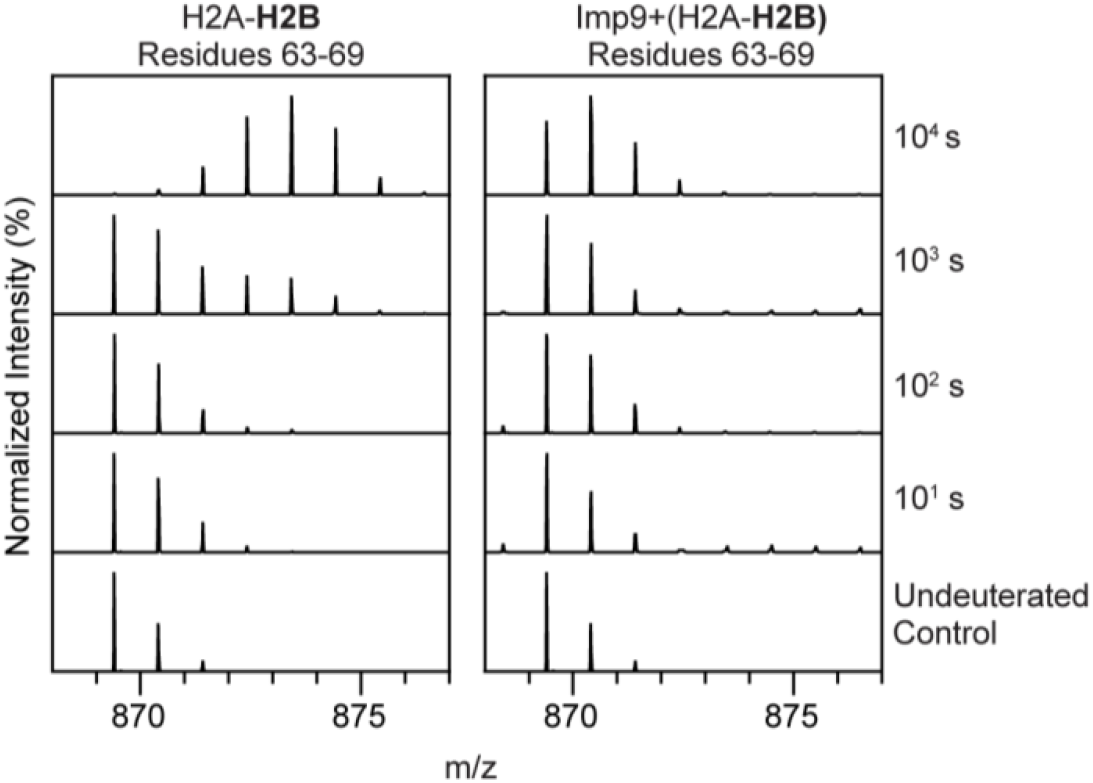
Spectra for H2B peptide, residues 63-69, in H2A-H2B (left) and Imp9+(H2A-H2B) (right). Spectra for the undeuterated control and 4 time points (10^1^,10^2^,10^3^, and 10^4^ s) are stacked. EX1 kinetics is most clearly seen in H2A-H2B alone at 10^3^s.

**Figure S5:**
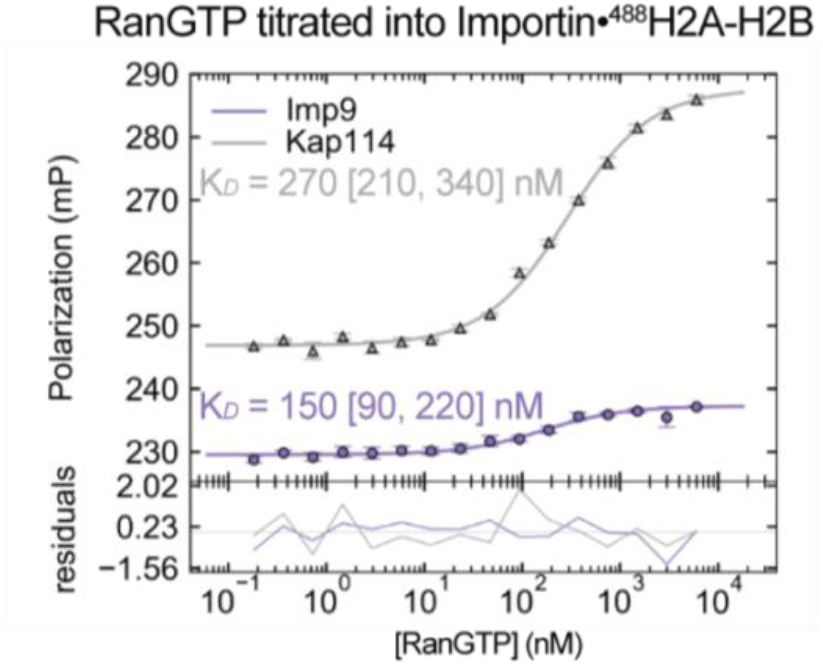
Affinity of RanGTP for Imp9/Kap114•H2A-H2B. Kap114 experiments are plotted in gray, and Imp9 experiments are plotted in light purple. RanGTP was titrated into Imp9/Kap114•^488^H2A-H2B. ^488^H2A-H2B contains H2A-K119C labeled with XFD488. The points in the upper plots are the mean of the triplicate measurements, and the error bars are ±1SD. The residual plots below show the deviation of the mean from the fitted line.

**Table S1:**
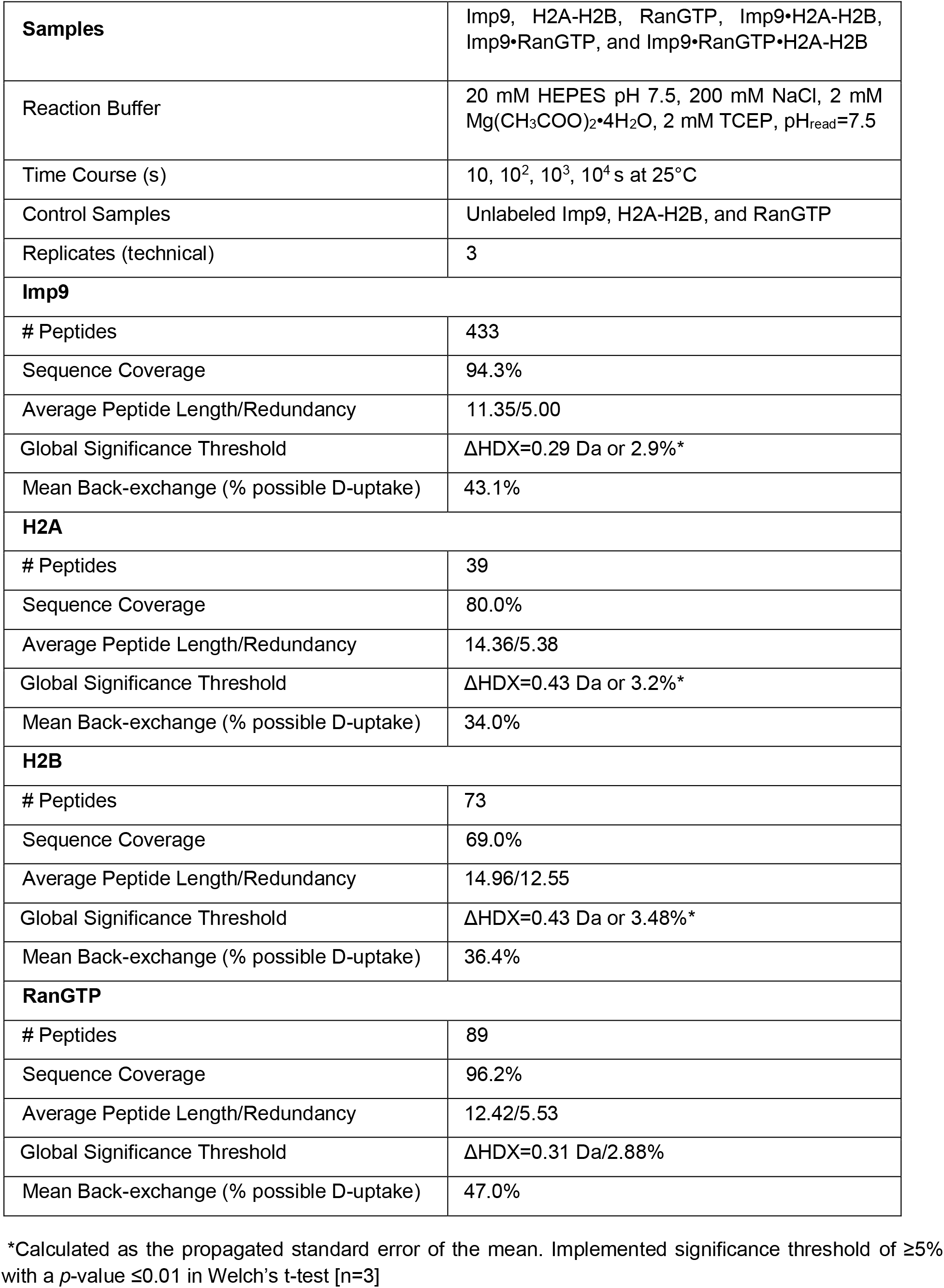
Summary of HDX data.

**Table S2: HDX state data for Imp9, H2A-H2B, and RanGTP (Attached as .csv)**

**Table S3:** HDX-based Prediction of the Affinity of RanGTP for Imp9 in the Ternary Complex.

| Region | Peptides | Average $\Delta\text{HDX} \pm \text{SD}$ Loss in Ternary Complex (%)<sup>*</sup> | Predicted $K_D$ (nM)<sup>**</sup> |
| --- | --- | --- | --- |
| Imp9 residues 38-52 that bind RanGTP | 38-45<br>38-46<br>39-46<br>42-51<br>46-51<br>46-52<br>47-52 | 45.3±10.8<br>[indicating only 54.7% of RanGTP bound to Imp9 in the ternary complex compared to 100% bound in the binary complex] | ~200 nM |

<sup>\*</sup>Calculated as $(1 - \frac{\text{Imp9} - \text{Imp9}_{\text{Ternary}}}{\text{Imp9} - \text{Imp9}_{\text{Binary}}}) \times 100\%$ , where $\text{Imp9}_{\text{Binary}}$ is Imp9•RanGTP at $10^4$ s.
<sup>\*\*</sup>Estimated using spreadsheet from Kochert et al. 2018 (32).

